# Light-sensitive phosphorylation regulates enzyme activity and filament assembly of human IMPDH1 retinal splice variants

**DOI:** 10.1101/2023.09.21.558867

**Authors:** S. John Calise, Audrey G. O’Neill, Anika L. Burrell, Miles S. Dickinson, Josephine Molfino, Charlie Clarke, Joel Quispe, David Sokolov, Rubén M. Buey, Justin M. Kollman

**Author notes:** corresponding author. Address: Box 357350, 1959 NE Pacific St, Seattle, WA 98195-7350. **Author contributions:** SJC, ALB, RMB, and JMK conceived the study; SJC, ALB, and JMK designed experiments; SJC, AGO, ALB, JM, CC, and RMB generated reagents; SJC, AGO, ALB, MSD, JM, CC, JQ, and DS performed experiments; SJC, AGO, and MSD analyzed data; JMK supervised and provided funding for the work; SJC wrote the original draft of the manuscript; SJC and JMK critically edited the manuscript. **Competing Interest Statement:** The authors declare that they have no competing interests.

## Abstract

Inosine monophosphate dehydrogenase (IMPDH) is the rate-limiting enzyme in *de novo* guanosine triphosphate (GTP) synthesis and is controlled by feedback inhibition and allosteric regulation. IMPDH assembles into micron-scale filaments in cells, which desensitizes the enzyme to feedback inhibition by GTP and boosts nucleotide production. The vertebrate retina expresses two tissue-specific splice variants IMPDH1(546) and IMPDH1(595). IMPDH1(546) filaments adopt high and low activity conformations, while IMPDH1(595) filaments maintain high activity. In bovine retinas, residue S477 is preferentially phosphorylated in the dark, but the effects on IMPDH1 activity and regulation are unclear. Here, we generated phosphomimetic mutants to investigate structural and functional consequences of phosphorylation in IMPDH1 variants. The S477D mutation re-sensitized both variants to GTP inhibition, but only blocked assembly of IMPDH1(595) filaments and not IMPDH1(546) filaments. Cryo-EM structures of both variants showed that S477D specifically blocks assembly of the high activity assembly interface, still allowing assembly of low activity IMPDH1(546) filaments. Finally, we discovered that S477D exerts a dominant-negative effect in cells, preventing endogenous IMPDH filament assembly. By modulating the structure and higher-order assembly of IMPDH, phosphorylation at S477 acts as a mechanism for downregulating retinal GTP synthesis in the dark, when nucleotide turnover is decreased. Like IMPDH1, many other metabolic enzymes dynamically assemble filamentous polymers that allosterically regulate activity. Our work suggests that posttranslational modifications may be yet another layer of regulatory control to finely tune activity by modulating filament assembly in response to changing metabolic demands.

**SIGNIFICANCE STATEMENT:** Over 20 different metabolic enzymes form micron-scale filaments in cells, suggesting that filament assembly is a conserved mechanism for regulating diverse metabolic pathways. Filament assembly regulates catalytic activity of many of these enzymes, including inosine monophosphate dehydrogenase (IMPDH), the rate-limiting enzyme in *de novo* GTP biosynthesis. The vertebrate retina expresses two IMPDH1 splice variants that are critical for maintaining nucleotide levels required for phototransduction. Here, we show that filament assembly by these variants is itself controlled by phosphorylation at a single residue, adding further complexity to the tight regulation of nucleotide metabolism in the retina. Phosphorylation and other posttranslational modifications are likely to be a general regulatory mechanism controlling filament assembly by enzymes in many different metabolic pathways.

## INTRODUCTION

Photoreceptor cells in the retina require high levels of guanine nucleotides to meet the metabolic demands of cyclic GMP (cGMP) signaling. To maintain nucleotide levels, they activate the *de novo* purine synthesis pathway to supplement nucleotides supplied by salvage pathways. Purine synthesis is a critical metabolic hub and enzymes in the pathway are under tight control by various modes of regulation, including feedback inhibition, allostery (Buey et al., 2015), biomolecular condensation (Pedley et al., 2022a), posttranslational modification (Liu et al., 2019), oligomerization and polymerization, and assembly of polymers into micron-scale subcellular structures known as metabolic filaments (Hvorecny and Kollman, 2023; Lynch et al., 2020; Park and Horton, 2019; Simonet et al., 2020).

The *de novo* purine synthesis pathway yields inosine monophosphate (IMP), the precursor for adenine and guanine nucleotides. IMP dehydrogenase (IMPDH) is the rate-limiting enzyme in guanosine triphosphate (GTP) synthesis (**Supp. Fig. 1A**). In humans, IMPDH has two canonical 514-residue isoforms with 84% sequence identity and similar kinetics (Carr et al., 1993). IMPDH1 is expressed at low levels in most tissues, while IMPDH2 is upregulated in proliferating and transformed cells (Collart et al., 1992; Jackson et al., 1975; Nagai et al., 1992; Senda and Natsumeda, 1994). IMPDH is regulated by GTP and ATP, which compete for three nucleotide binding sites in the regulatory Bateman domain to inhibit or activate the enzyme, respectively (Buey et al., 2022). In solution, IMPDH forms a catalytic tetramer that reversibly dimerizes upon nucleotide binding to form a functional octamer, which can assemble end-on-end to form a polymer (referred to as “filaments” here). GTP binds nucleotide binding sites 2 and 3, while ATP binds site 1 and competes with GTP for site 2 (**Supp. Fig. 1B**) (Buey et al., 2015; Buey et al., 2017; Johnson and Kollman, 2020).

Binding of GTP and ATP controls the polymerization of IMPDH *in vitro*. The presence of ATP in sites 1 and 2 promotes filaments composed of octamers in an extended, high activity state, while GTP in sites 2 and 3 leads to filaments containing compressed, less active octamers (**Supp. Fig. 1C,D**) (Anthony et al., 2017; Burrell et al., 2022; Johnson and Kollman, 2020). For IMPDH2, filament assembly maintains a “flat” conformation of the catalytic tetramer that prevents full inhibition of the enzyme within the filament, allowing the enzyme to remain partially active *in vitro* (Fernández-Justel et al., 2019; Johnson and Kollman, 2020). In cells, single IMPDH filaments bundle together to form micron-scale ultrastructures (Juda et al., 2014; Schiavon et al., 2018; Thomas et al., 2012). IMPDH filament assembly *in vivo* correlates with metabolic conditions that require high GTP levels, like T cell activation and the rapid proliferation of stem cells (Calise and Chan, 2020; Calise et al., 2018; Carcamo et al., 2011; Duong-Ly et al., 2018; Keppeke et al., 2018).

IMPDH1 is the dominant isoform in the vertebrate retina and is expressed as two tissue-specific splice variants (Bowne et al., 2006b; Hedstrom, 2009). IMPDH1(546) has a disordered extension of 37 residues that replaces the five canonical residues at the C-terminus. IMPDH1(595) contains the same C-terminal extension plus 49 residues at the N-terminus (**Supp. Fig. 2A**) (Burrell and Kollman, 2022; Spellicy et al., 2007). IMPDH1(546) and IMPDH1(595) are less sensitive to GTP inhibition compared to the canonical enzyme IMPDH1(514), enabling increased GTP production to meet the significant demands of cGMP signaling in photoreceptor cells (**Supp. Fig. 2B**) (Andashti et al., 2020; Andashti et al., 2021; Burrell et al., 2022; Plana-Bonamaisó et al., 2020). Within IMPDH1(514) and IMPDH1(546) filaments, the assembly interface between two octamers can adopt two conformations referred to as the large and small interfaces, which correlate with high and low enzymatic activity, respectively (Burrell et al., 2022). For IMPDH1(595), the N-terminal extension locks it in the large interface, which stabilizes a more active conformation of the enzyme and contributes to increased resistance to GTP inhibition compared to the canonical enzyme (**Supp. Fig. 2C**) (Burrell et al., 2022).

The importance of IMPDH1 in retinal function is reflected in the multiple missense mutations that have been linked to gradual vision loss caused by autosomal dominant retinitis pigmentosa (Bowne et al., 2002; Bowne et al., 2006a; Grover et al., 2004; Kennan et al., 2002; Wada et al., 2005) and Leber congenital amaurosis (Bowne et al., 2006a). None of the mutations alter canonical IMPDH1 catalytic activity *in vitro* (Aherne et al., 2004; Mortimer and Hedstrom, 2005; Xu et al., 2008), but we recently showed that some mutations alter the architecture of IMPDH retinal variant filaments, disrupting regulation by GTP and providing clues as to why clinical manifestations are limited to the retina (Burrell et al., 2022).

Recent phosphoproteomic analyses of bovine retinas identified three phosphorylation sites in IMPDH1 associated with light and dark states (Plana-Bonamaisó et al., 2020). Phosphorylation occurs at T159/S160, residues directly involved with allosteric nucleotide binding site 1, in response to light. This desensitizes the enzyme to GTP regulation, allowing elevated GTP pools for phototransduction. Phosphorylation at S416, a residue on a mobile flap in the catalytic domain required for catalysis, was observed in both light and dark states. The third site S477 is preferentially phosphorylated in the dark, but its effects on reaction kinetics and GTP regulation are unclear. An S477D phosphomimetic mutant of IMPDH1(546) had similar reaction kinetics to wildtype in the absence of nucleotides, and GTP regulation of IMPDH1(514)-S477D appeared similar to wildtype canonical enzyme, although no data on the GTP inhibition of S477D mutant retinal variants were shown (Plana-Bonamaisó et al., 2020). Phosphorylation at S477 was speculated to disrupt filament formation (Plana-Bonamaisó et al., 2020).

Here, we show that S477D disrupts assembly of the large, high activity filament interface, completely preventing IMPDH1(595) filament assembly and forcing IMPDH1(546) filaments into the small, lower activity interface. These changes in higher-order structure correspond to increased sensitivity to GTP inhibition and lower reaction velocity in enzyme assays. This suggests that phosphorylation directly affects flux through IMPDH1 by preventing its ability to form high-activity filaments, introducing an additional mechanism for controlling enzyme filament formation to tune metabolic activity.

## RESULTS

### S477D disrupts IMPDH1(595) filament assembly *in vitro*

We examined purified S477D variants IMPDH1(595)-S477D and IMPDH1(546)-S477D by negative stain electron microscopy (EM) in the presence of no ligand (Apo) or either 1 mM ATP or 1 mM GTP, which promote polymerization of the wildtype enzyme (Burrell et al., 2022; Johnson and Kollman, 2020). S477D completely prevented assembly of IMPDH1(595) filaments, allowing only free octamer formation in the presence of ATP or GTP (**Fig. 1A**). At this resolution, higher-order assembly of IMPDH1(546)-S477D appeared unaffected, assembling octamers and polymers similar to wildtype enzyme (**Fig. 1B**). S477D also partially prevented polymerization of the canonical IMPDH1(514), although the effects were not as clear as in the retinal variants (**Supp. Fig. 3**). Previous structures of IMPDH1 filaments showed that S477 contacts the N-terminus of the opposing monomer in the large interface, but makes no contacts in the small interface, suggesting phosphorylation at S477 is likely to specifically break the large interface (**Supp. Fig. 2D**). Thus, we suspected that the IMPDH1(546)-S477D filaments we observed by negative stain were in the small interface. Complete disassembly of IMPDH1(595) filaments supports our previous structural data showing that IMPDH1(595) filament assembly is restricted to the large interface (Burrell et al., 2022).

**Figure 1.**
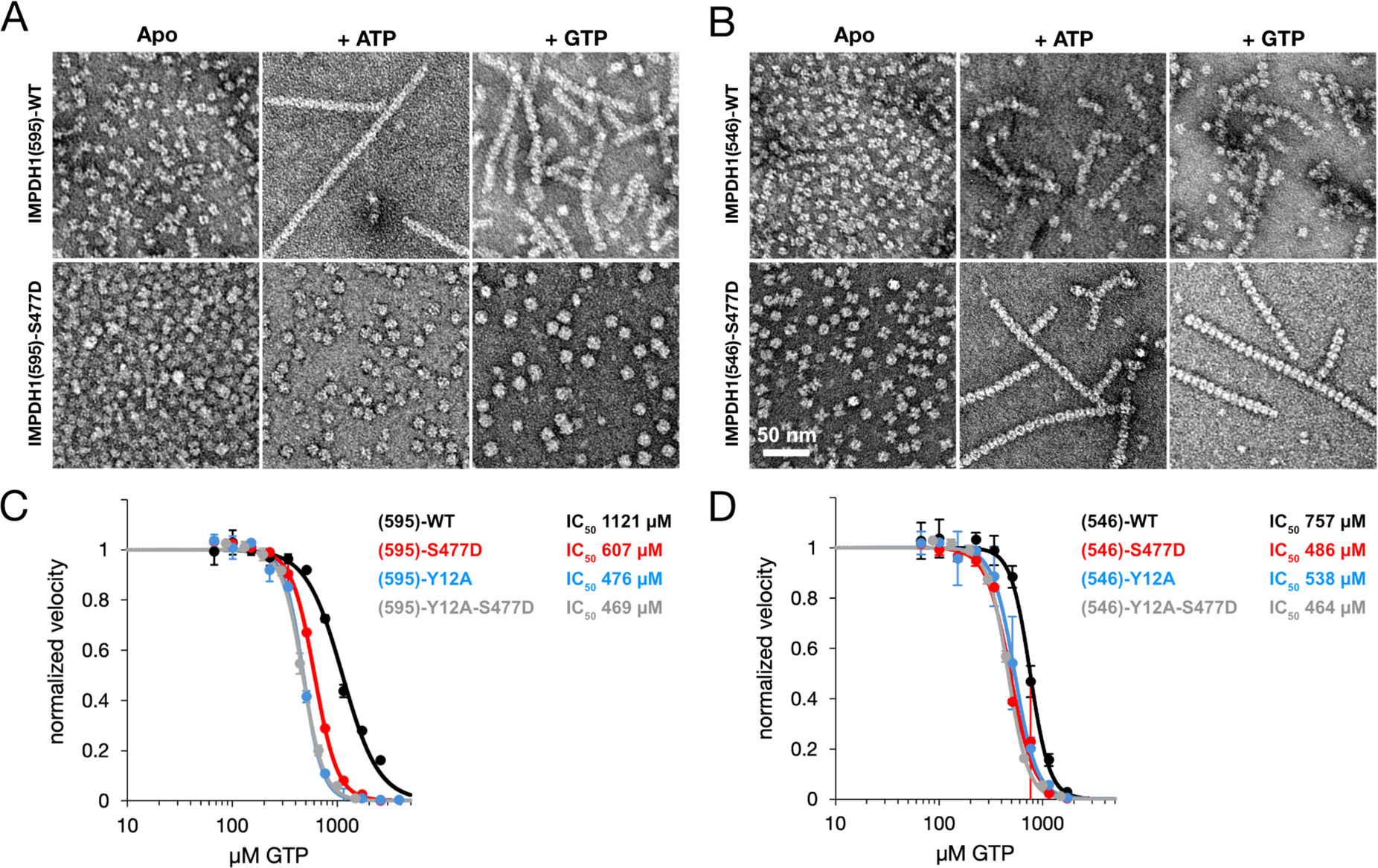
S477D disrupts IMPDH1(595) filament assembly and re-sensitizes both variants to GTP inhibition. **(A)** Negative stain EM of IMPDH1(595)-WT and IMPDH1(595)-S477D under Apo, + 1 mM ATP, or + 1 mM GTP conditions. **(B)** Negative stain EM of IMPDH1(546)-WT and IMPDH1(546)-S477D under the same conditions. **(C)** GTP inhibition curve of IMPDH1(595)-S477D (red) compared to WT (black), Y12A (blue), and Y12A-S477D (grey). **(D)** GTP inhibition curve of IMPDH1(546)-S477D (red) compared to WT (black), Y12A (blue), and Y12A-S477D (grey).

### S477D re-sensitizes both retinal variants to GTP inhibition

Wildtype (WT) retinal variants IMPDH1(546)-WT and IMPDH1(595)-WT are significantly less sensitive to GTP inhibition compared to the canonical IMPDH1(514) (Burrell et al., 2022). Here, we wanted to test the S477D mutants for any effects on GTP regulation of activity. In both retinal variants, S477D re-sensitized the enzyme to GTP inhibition. S477D led to a two-fold decrease in the IC_50_ for GTP in IMPDH1(595) (**Fig. 1C**) and a 1.5-fold decrease in IMPDH1(546) (**Fig. 1D**). In both cases, the IC_50_ for GTP was reduced to the same magnitude observed in the engineered mutation Y12A, which disrupts the filament assembly interface (Anthony et al., 2017; Burrell et al., 2022; Fernández-Justel et al., 2019; Johnson and Kollman, 2020).

Inhibition of IMPDH1 involves two conformational changes: interdomain movements that switch from an extended to compressed octamer, and transition of the catalytic tetramer from a flat conformation to a bowed conformation at the filament assembly interface (Burrell et al., 2022; Johnson and Kollman, 2020). The large interface is associated with stabilizing the flat conformation, leading to higher catalytic activity, while the small interface is associated with the bowed conformation and lower activity. In completely breaking IMPDH1(595) filament assembly, S477D removes the enzyme’s ability to engage the large interface and stabilize the flat conformation. Free IMPDH octamers adopt the compressed and bowed conformation upon GTP binding, leading to lower activity (Burrell et al., 2022; Johnson and Kollman, 2020), explaining the increased sensitivity to GTP inhibition of IMPDH1(595)-S477D in our assays. However, why IMPDH1(546)-S477D was re-sensitized to the levels of the engineered filament-disrupting mutation Y12A was not clear. To gain further insights into structural details, we turned to cryogenic EM (cryo-EM).

### IMPDH1(595)-S477D free octamer adopts a bowed conformation

First, we solved a 3.1 Å cryo-EM structure of the IMPDH1(595)-S477D free octamer in the presence of GTP, ATP, IMP, and NAD^+^ (**Fig. 2A, Supp. Fig. 4**). We built an atomic model and aligned it with a previous model (PDB: 7RGD) of IMPDH1(595)-WT under the same ligand conditions, in a compressed state within a filament. We observed no significant changes in the conformation of the backbone, with a calculated root-mean-square deviation (RMSD) of 0.569 Å. We then took a closer look at the catalytic tetramer of each model to determine if IMPDH1(595)-S477D was in the flat or bowed conformation (**Fig. 2B**). We aligned S477D and WT tetramers on chain A and observed a 3° shift in chain C of S477D relative to WT, showing that IMPDH1(595)-S477D free octamer adopts a bowed conformation similar to the inhibited canonical IMPDH1(514)-WT within a filament (RMSD IMPDH1(595)-S477D vs. inhibited IMPDH1(514)-WT = 0.684 Å). IMPDH1(595)-WT is stabilized in the large interface and flat conformation in the filament by its extended N-terminus (Burrell et al., 2022). The extended N-terminus was unresolved in our map of the free octamer, suggesting it only becomes ordered upon interaction with a binding partner.

**Figure 2.**
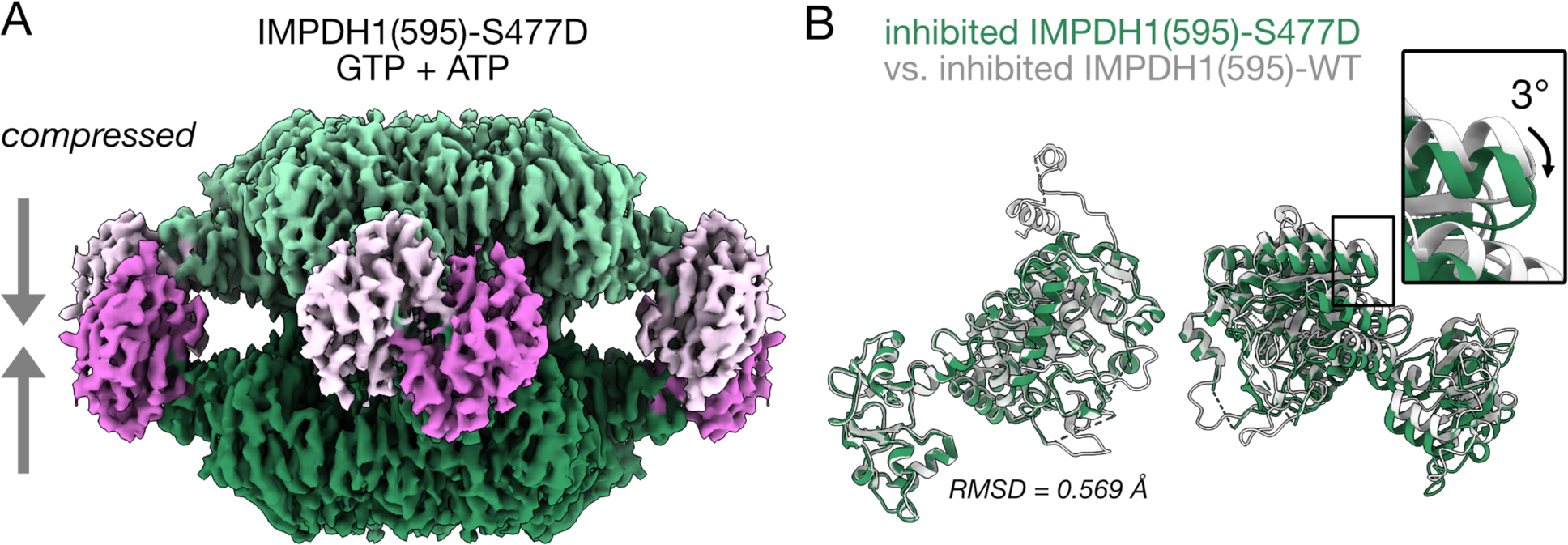
Cryo-EM structure of the IMPDH1(595)-S477D free octamer. **(A)** Cryo-EM map (3.1 Å) of the compressed IMPDH1(595)-S477D free octamer under inhibitory conditions, in the presence of GTP, ATP, IMP, and NAD^+^. **(B)** Alignment of the atomic model built from the map in panel A (green) with previously built model of IMPDH1(595)-WT (grey, PDB: 7RGD). The S477D catalytic tetramer is in a bowed conformation, bent approximately 3° relative to the flat conformation of the WT.

### Both inhibited and active IMPDH1(546)-S477D filaments assemble with the small interface

Next, we solved a 3.3 Å cryo-EM structure of the IMPDH1(546)-S477D compressed octamer within a filament in the presence of GTP, ATP, IMP, and NAD^+^ (**Fig. 3A, Supp. Fig. 5**). We built an atomic model of the compressed octamer and compared it to a previous model (PDB: 7RGQ) of IMPDH1(546)-WT in a compressed state within a filament, which adopts the small interface (Burrell et al., 2022). Aligning these two models showed no significant differences between single protomers or octamers (RMSD = 0.534 Å). From the same data set we separately determined a structure of the filament assembly interface by centering the reconstruction on the D4 symmetry center between octamers in the filament, an approach that we have used in the past to yield the highest resolution reconstructions of the interface (Burrell et al. 2022, Johnson and Kollman, 2020). This approach yielded a 3.3 Å structure of the assembly contacts, which is essentially identical to the IMPDH1(546)-WT assembly interface (RMSD = 0.589 Å) (**Fig. 3C; Supp. Fig. 6**). Thus, at the level of protomer, octamer, and filament the S477D mutation does not affect the structure of fully inhibited IMDPH1(546).

**Figure 3.**
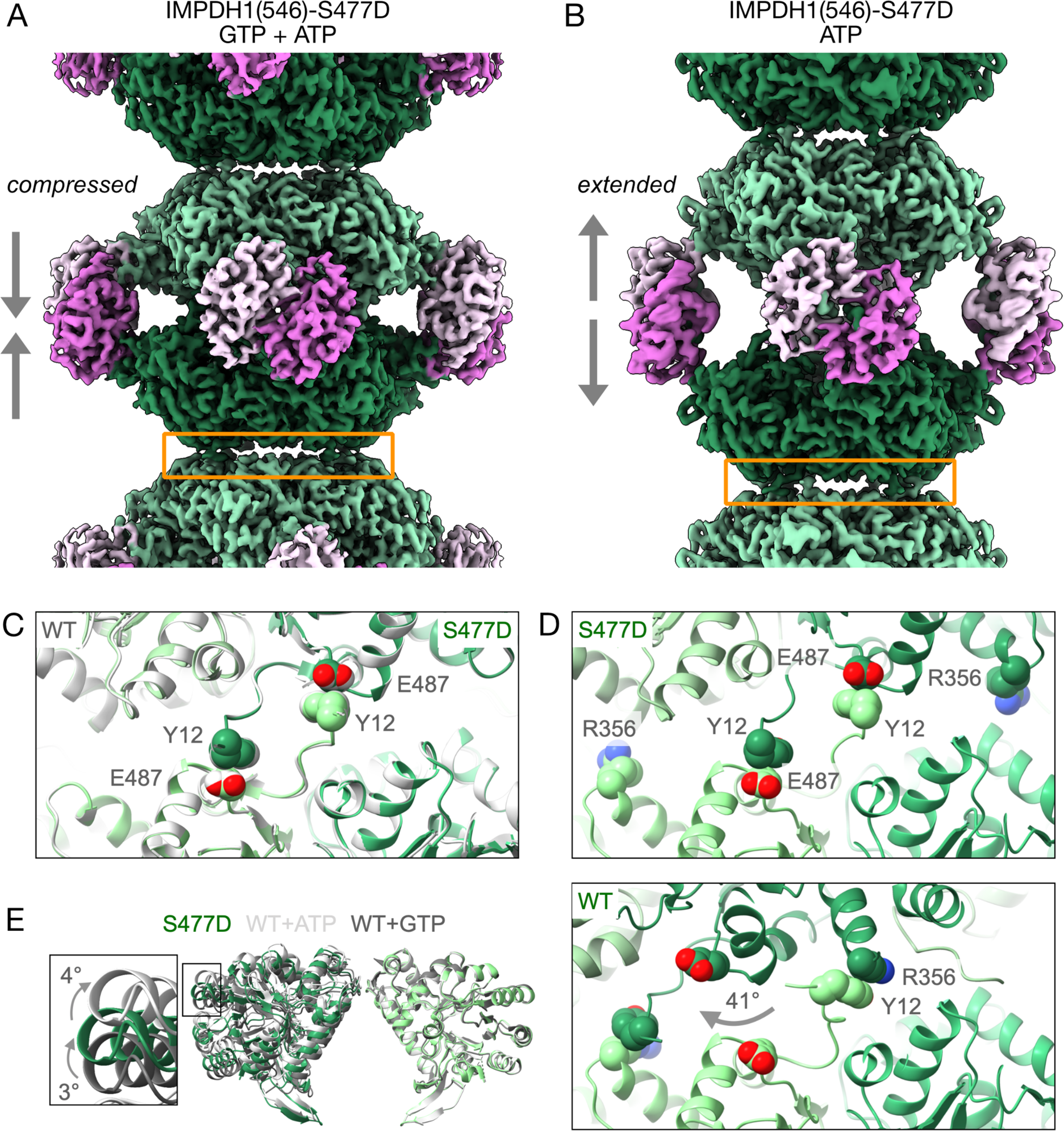
Cryo-EM structures of IMPDH1(546)-S477D in inhibited and active states. **(A)** Cryo-EM map (3.3 Å) of a compressed IMPDH1(546)-S477D octamer within a filament, in the presence of GTP, ATP, IMP, and NAD^+^. Orange box: assembly interface shown in panel C. **(B)** Cryo-EM map (2.4 Å) of an extended IMPDH1(546)-S477D octamer within a filament, in the presence of ATP, IMP, and NAD^+^. Orange box: assembly interface shown in panel D. **(C)** Assembly interface of the compressed IMPDH1(546)-S477D filament (green) aligned to compressed IMPDH1(546)-WT (grey, PDB: 7RGI). In both S477D and WT, the interaction between Y12 and E487 drives filament assembly. **(D)** Extended IMPDH1(546)-S477D filament assembly interface (top) compared to the extended IMPDH1(546)-WT interface (bottom, PDB: 7RGL). Top: like the compressed 546 filament, Y12 and E487 drive assembly of the extended 546 filament. Bottom: in the extended 546-WT filament, Y12 and R356 interact to drive assembly. **(E)** Active IMPDH1(546)-WT (light grey), active IMPDH1(546)-S477D (green), and inactive IMPDH1(546)-WT (dark grey) aligned on the right monomer. S477D adopts an intermediate conformation between the fully bowed and flat conformations.

In the active state in the presence of ATP, IMP, and NAD^+^, IMPDH1(546)-WT assembles a filament with the large interface (Burrell et al., 2022). We solved a 2.4 Å cryo-EM structure of IMPDH1(546)-S477D under the same conditions (**Fig. 3B, Supp. Fig. 7**) and aligned the model to IMPDH1(546)-WT in an extended state within a filament (PDB: 7RGM). Individual protomers appeared highly similar (RMSD = 0.557 Å) but when we aligned octamers on chain A, we detected a 3° shift in the protomer on the opposite side of the octamer. We solved a 2.1 Å interface-centered structure of IMPDH1(546)-S477D and compared the filament assembly interface to IMPDH1(546)-WT (**Supp. Fig. 8**). The IMPDH1(546)-S477D extended, active filament adopts the small interface driven by contacts between Y12 and E487 (**Fig. 3D, top**). This is significantly different from the IMPDH1(546)-WT large interface, which has a relative 41° rotation with R356 contacting Y12 (**Fig. 3D, bottom**). We then aligned active IMPDH1(546)-WT, active IMPDH1(546)-S477D, and inhibited IMPDH1(546)-S477D to compare their catalytic tetramer conformations. The catalytic tetramer of active IMPDH1(546)-S477D adopts a unique intermediate conformation between bowed and flat (**Fig. 3E, inset**). Forming the small interface with an intermediate tetramer conformation suggests the enzyme may be primed to enter the compressed, bent state when GTP is present, lowering the transition barrier and effectively raising the affinity for GTP. In a kinetics assay, this would mean IMPDH1(546)-S477D is more sensitive to GTP inhibition, as we detected in **Fig. 1D**.

### S477D is a dominant-negative regulator of IMPDH filament assembly

Given the obvious disruption to higher-order assembly of IMPDH1(595)-S477D, we wanted to investigate the impact of the phosphomimetic on filament assembly in cells. We transfected HeLa cells with expression constructs encoding FLAG-tagged WT, S477D mutant, and S477A mutant retinal variants (**Fig. 4**). After 48 hours, we treated cells with 1 mM ribavirin for 2 hours, which leads to robust IMPDH filament assembly in nearly 100% of cells (Calise et al., 2016; Carcamo et al., 2011; Covini et al., 2012; Keppeke et al., 2015). We then fixed and stained cells with an anti-IMPDH2 antibody to label endogenous IMPDH and anti-FLAG antibody to label our transfected protein.

**Figure 4.**
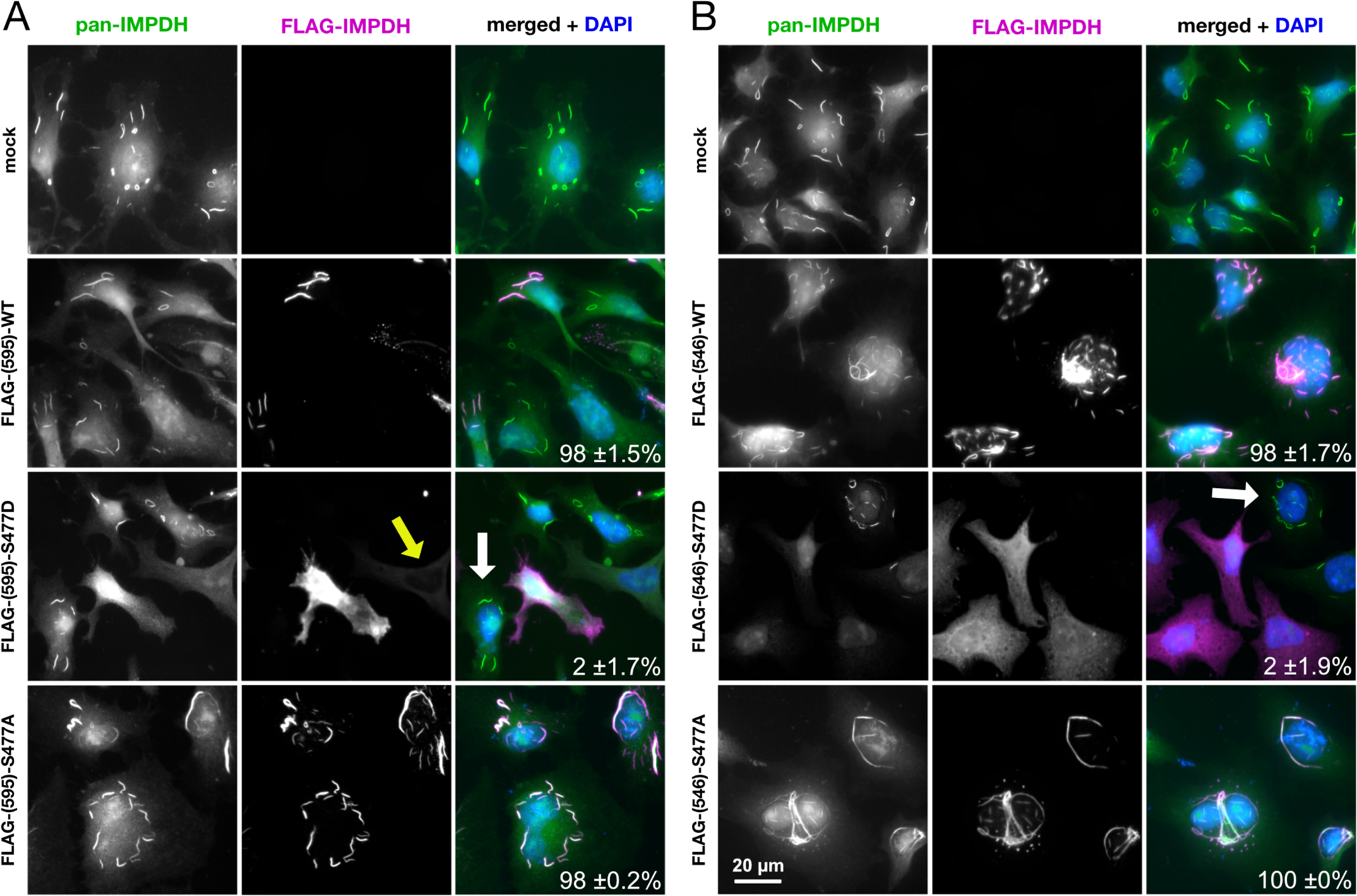
S477D disrupts filament assembly in a dominant-negative manner in living cells. **(A)** HeLa cells were transfected with FLAG-tagged IMPDH1(595)-WT, IMPDH1(595)-S477D, or IMPDH1(595)-S477A, then treated with 1 mM ribavirin for 2 h to induce filament assembly. Mock transfection is shown as a control. Green: all IMPDH (endogenous + transfected). Magenta: FLAG-tagged fusion proteins. Blue: nuclei stained with DAPI. White arrow: non-transfected cell with filament formation, serving as an internal control. Yellow arrow: cell with low overexpression of FLAG-IMPDH1(595)-S477D. Percentage of transfected cells containing filaments is displayed in the bottom right corner of merged panels. **(B)** Same experiment as in panel A, but with FLAG-tagged IMPDH1(546)-WT, IMPDH1(546)-S477D, and IMPDH1(546)-S477A. Experiments were performed twice.

Both WT and S477A variants assembled typical micron-scale filaments in a high percentage of transfected cells, as expected (595-WT: 98%, 595-S477A: 98%, 546-WT: 98%, 546-S477A: 100%). However, S477D exerted a dominant-negative effect, blocking essentially all filament assembly by endogenous IMPDH in transfected cells (595-S477D: 2%, 546-S477D: 2%). The dominant-negative effect was not due to increased overexpression of the S477D variants relative to WT and S477A, as cells with low S477D overexpression also completely lacked filaments (yellow arrow, **Fig. 4A**). In cell populations transfected with S477D mutants, filaments can be observed in adjacent non-transfected cells as an internal control (white arrows, **Fig. 4**). Since overexpression alone also causes IMPDH1 filament assembly (Gunter et al., 2008; Keppeke et al., 2018; Keppeke et al., 2023), we performed the same experiment without ribavirin treatment and observed similar results (**Supp. Fig. 9**).

These data suggest two important points: (1) human IMPDH1 and IMPDH2 co-assemble into the same filament (Keppeke et al., 2023) and (2) because IMPDH1(546)-S477D forms filaments *in vitro*, its inability to form filaments in cells suggests that IMPDH filaments in living cells predominantly assemble using the large interface. Together, these points imply that phosphorylation at S477 is a molecular mechanism for regulating flux through *de novo* GTP synthesis by controlling higher-order assembly of the pathway’s rate-limiting enzyme.

## DISCUSSION

Despite the increasing interest in higher-order assembly of IMPDH, posttranslational modifications of the enzyme have received little attention. However, a recent phosphoproteomic analysis of bovine retinas adapted to light and dark states shed light on three novel phosphorylation sites in IMPDH1: T159/S160, S416, and S477 (Plana-Bonamaisó et al., 2020). The study focused mainly on light-dependent phosphorylation at T159/S160, which desensitizes IMPDH1 to GTP inhibition to boost GTP levels in support of phototransduction. On the other hand, phosphorylation at S477 was correlated with the dark, when less GTP production is needed (Plana-Bonamaisó et al., 2020). S477D seemed to have no effect on basal IMPDH1(546) activity *in vitro*, suggesting that S477D might instead play a structural role in disrupting filament assembly.

Here, we showed that S477D re-sensitizes human retinal IMPDH1 variants to GTP inhibition and prevents filament assembly in living cells in a dominant-negative manner, which would lead to overall decrease in cellular GTP pools. Our data demonstrate a mechanism for control of IMPDH filament assembly dynamics and GTP production by phosphorylation at a single site. We propose that phosphorylation at S477 reduces the extent of IMPDH filaments in photoreceptor cells in the dark, leading to a global decrease in GTP levels when nucleotide turnover rate is reduced. In the light, an unknown phosphatase removes the modification, allowing filaments to reassemble and boost GTP levels to support cGMP signaling again (**Fig. 5**).

**Figure 5.**
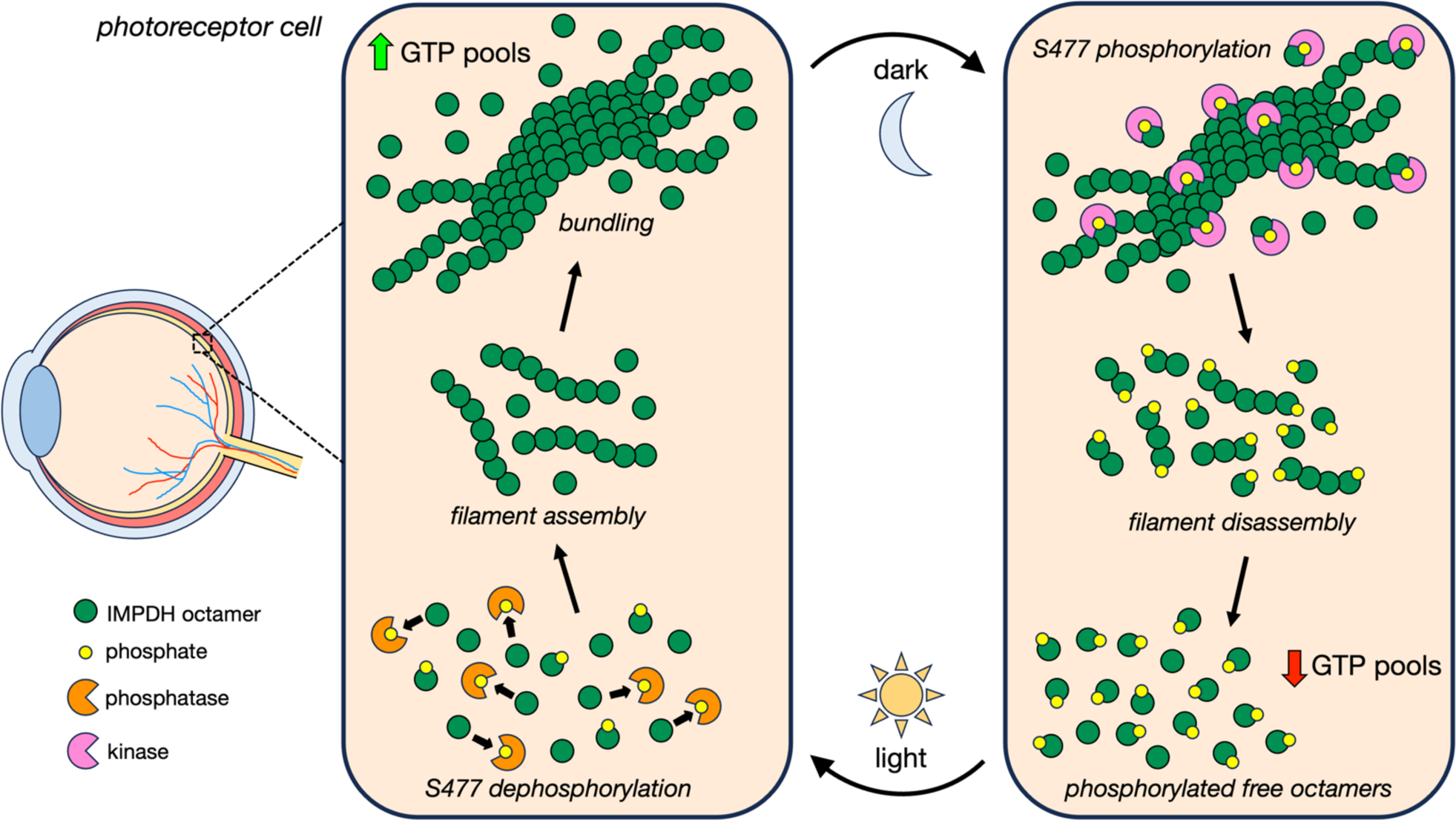
Model of IMPDH1-S477 phosphorylation in the retina. In the dark, phosphorylation of IMPDH1 at S477 disassembles filaments and the cellular pool of IMPDH shifts towards phosphorylated free octamers, leading to a decrease in GTP pools when less nucleotide turnover is occurring. In the light, S477 is dephosphorylated and filaments reassemble to boost GTP production when nucleotide turnover is higher. The cycle of IMPDH1 filament assembly and disassembly in light and dark states continues to regulate cellular GTP levels.

Given our *in vitro* data, we were surprised that IMPDH1(546)-S477D failed to form filaments in cells. The dominant-negative effect of S477D suggests that IMPDH1(546)-S477D co-assembles with endogenous IMPDH2 into filaments. Structural data show IMPDH2 forms filaments exclusively with the large interface (Burrell et al., 2022; Johnson and Kollman, 2020). Thus, in our experiments, co-assembly of IMPDH2 and IMPDH1(546)-S477D at the tetramer or octamer level would produce incompatible filament assembly interfaces. In the retina, the co-assembly of IMPDH1(595) (also restricted to the large interface) with IMPDH1(546) would have the same effect, and IMPDH1(546) would only exist in filaments with the large interface. Further studies in genome-edited cells expressing only one IMPDH variant at a time may provide further insight into *in vivo* assembly of the different forms of IMPDH.

We recently characterized IMPDH1 filaments in the zebrafish retina, finding a circadian pattern to filament assembly, with elongated filaments forming in photoreceptors during the day and smaller punctate structures at night (Cleghorn et al., 2022). We also showed that the zebrafish IMPDH1 retinal variants are very similar to human IMPDH1 in terms of structure, filament assembly, and regulation (Cleghorn et al., 2022). This is consistent with our finding here that the S477D mutation, which mimics dark-dependent phosphorylation, disrupts filament assembly and reduces enzyme activity in the presence of GTP. Besides zebrafish, IMPDH filament assembly has also been observed in mouse retinas, although the functional significance has yet to be described (Fernández-Justel et al., 2019).

Studies in cultured cells and mice have demonstrated a strong link between IMPDH filament assembly and cells that consume high levels of GTP. Robust IMPDH filament assembly occurs in stem cells (Carcamo et al., 2011; Keppeke et al., 2018) and lymphocytes (Calise et al., 2018; Duong-Ly et al., 2018), which require increased nucleotides to support rapid proliferation (Dayton et al., 1994; Fairbanks et al., 1995; Quéméneur et al., 2003). Polymerization of IMPDH1(546), IMPDH1(595), and IMPDH2 partially prevents feedback inhibition by guanine nucleotides compared to the unpolymerized state (Burrell et al., 2022; Fernández-Justel et al., 2019; Johnson and Kollman, 2020). In cells, this would allow accumulation of GTP to higher steady state levels, supporting rapid proliferation (IMPDH2) or phototransduction in photoreceptor cells (IMPDH1 retinal variants).

More than 20 different metabolic enzymes form filaments in cells, suggesting that filament assembly is a general mechanism for regulating multiple metabolic pathways (Hvorecny and Kollman, 2023; Lynch et al., 2020; Narayanaswamy et al., 2009; Noree et al., 2010; Park and Horton, 2019; Park and Horton, 2020; Shen et al., 2016; Simonet et al., 2020). For many metabolic enzymes, filament formation is an important mechanism for fine-tuning catalytic activity (Barry et al., 2014; Hansen et al., 2021; Hu et al., 2022; Hunkeler et al., 2018; Hvorecny et al., 2023; Lu et al., 2023; Lynch and Kollman, 2020; Lynch et al., 2017; Polley et al., 2019; Stoddard et al., 2020; Zhong et al., 2022). Here, we demonstrate that filament assembly itself can be directly tuned by phosphorylation, adding an additional layer to an already complex mode of regulation for IMPDH that occurs at different levels. Activity of IMPDH tetramers is modulated by feedback inhibition and oligomerization into octamers. IMPDH octamers are then regulated further by allosteric binding of nucleotides to the Bateman domain, which compresses or extends the conformation of the octamer. Polymerization of octamers into filaments can then modulate sensitivity to GTP inhibition. Filament assembly itself is then regulated by phosphorylation at S477 in response to changing metabolic conditions of the cell.

IMPDH localizes to the purinosome (Zhao et al., 2015), a biomolecular condensate assembled by the six enzymes directly upstream of IMPDH in *de novo* purine synthesis (An et al., 2008; Pedley et al., 2022b). Purinosome assembly correlates with increased flux through the pathway (Pareek et al., 2020; Zhao et al., 2015) and has been hypothesized to be regulated by posttranslational modifications (Liu et al., 2019). In fact, purinosome assembly is modulated by casein kinase II, AMP-activated protein kinase, and 3-phosphoinositide-dependent protein kinase 1, suggesting various signaling cascades spatially regulate purinosome enzymes (An et al., 2010; Schmitt et al., 2016; Schmitt et al., 2018). It is enticing to speculate that a network of phosphorylation and/or other modifications may spatiotemporally coordinate assembly of IMPDH into various subcellular compartments depending on the metabolic state of the cell.

Given the widespread utilization of filament assembly as a regulatory mechanism, it is likely that other polymerizing enzymes are regulated by posttranslational modification in the same way. Some correlations between posttranslational modification and IMPDH or cytidine triphosphate synthetase (CTPS) filament assembly in cells have been reported. Ankyrin repeat domain-containing protein 9 (ANKRD9) is a component of a cullin-RING superfamily E3 ligase complex that ubiquitinates both isoforms of IMPDH for degradation (Lee et al., 2018). ANKRD9 also localizes to and stabilizes IMPDH filaments induced by serum starvation in HeLa cells, suggesting ubiquitination plays some role in filament assembly (Hayward et al., 2019). The E3 ubiquitin ligase activity of Cbl is required for CTPS filament assembly in *Drosophila*, although it is not clear if the ubiquitination is direct (Wang et al., 2015). In a similar fashion, protein methylation is required for CTPS filament assembly in human cells (Lin et al., 2018). The mechanisms underlying these modifications remain unclear, but our current study opens the possibility for direct posttranslational modification as a general mechanism for modulating filament assembly dynamics.

The data presented here establish a molecular mechanism for reversible phosphorylation controlling assembly of IMPDH filaments and regulating GTP levels in the retina. This mechanism is likely to apply to a variety of filament-forming enzymes coordinating many different metabolic pathways in the cell. Future studies will determine if phosphorylation or other posttranslational modifications are the mechanism of control underlying general metabolic filament assembly *in vivo*.

## MATERIALS AND METHODS

### Molecular cloning

Kanamycin-resistant pSMT3 bacterial expression vectors containing coding sequences for human IMPDH1(546) and IMPDH1(595) with N-terminal 6xHis-SUMO tags were generated previously (Burrell et al., 2022). To generate S477D mutant versions, overlapping primers 5′-CGCAGCCTG<u>GA</u>TGTCCTTCGGTCCA-3′ and 5′-TGGACCGAAGGACA<u>TC</u>CAGGCTGCG-3′ containing a TC>GA change to replace a serine with an aspartic acid codon were used to amplify each plasmid. Plasmids were ligated at 50 °C for 1 hour in a Gibson assembly reaction. The reaction product was treated with DpnI enzyme at 37 °C for 30 min to degrade template DNA, then used for transformation of competent TOP10 *E. coli* cells. Transformed cells were cultured overnight on LB agar plates containing 50 µg/mL kanamycin. Single colonies were picked and cultured overnight in LB medium + 50 µg/mL kanamycin and recombinant plasmids isolated using the GeneJET Plasmid Miniprep Kit (Thermo Fisher #K0503). Insert sequences in each plasmid were confirmed by Sanger sequencing. Vector sequence was previously confirmed by Sanger sequencing of parental constructs.

For mammalian expression, coding sequences for wildtype and S477D retinal variants were subcloned from the above constructs into a pCMV3 vector (generously provided by Dr. Gerson Keppeke, Universidad Católica del Norte, Coquimbo, Chile). S477D constructs were then used as templates for PCR reactions to generate full-length S477A constructs, which were then self-ligated by Gibson assembly. Primers used for cloning are listed in **Supp. Table 1**. The same procedure was used for cloning mammalian constructs as bacterial expression constructs (see above), except that different primers were used. Full-length plasmid sequencing was performed on all mammalian constructs by Nanopore sequencing. Insert regions were also validated by Sanger sequencing.

### Recombinant IMPDH1 expression and purification

Recombinant human IMPDH1 retinal variant proteins were expressed and purified as described previously (Anthony et al., 2017; Burrell et al., 2022; Cleghorn et al., 2022). Briefly, BL21 (DE3) *E. coli* cells were transformed with wildtype or S477D mutant IMPDH1(546) or IMPDH1(595) expression vectors. Cells were cultured in LB medium at 37 °C until reaching an optical density (OD_600_) of 0.9, then expression was induced by addition of isopropyl-β-d-thiogalactoside (IPTG) to 1 mM for 4 h at 30 °C. Cells were pelleted and flash frozen in liquid nitrogen and stored at −80 °C until use.

Pellets were thawed, resuspended in lysis buffer (50 mM KPO_4_, 300 mM KCl, 10 mM imidazole, 800 mM urea, pH 8.0), and lysed with an Emulsiflex-05 homogenizer. Lysate was clarified by centrifugation at 33,746 x g for 30 min at 4 °C and 6xHis-SUMO-tagged IMPDH1 was purified by immobilized metal ion affinity chromatography using HisTrap FF nickel sepharose columns (Cytiva) on an ӒKTA Start chromatography system. After on-column washing with lysis buffer, protein was eluted in buffer (50 mM KPO_4_, 300 mM KCl, 500 mM imidazole, pH 8.0) and fractionated into 1 mL volumes. Peak fractions were combined and incubated with 1 mg ULP1 protease (Malakhov et al., 2004) per 100 mg IMPDH1 for 1 h on ice, then supplemented with 1 mM dithiothreitol (DTT) and 800 mM urea. Protein was concentrated with a 30,000 molecular weight cutoff filter (Amicon) and size-exclusion chromatography was performed on an ӒKTA Pure chromatography system using a Superose 6 column (Cytiva) pre-equilibrated in gel filtration buffer (20 mM HEPES, 100 mM KCl, 800 mM urea, 1 mM DTT, pH 8.0). Peak fractions were concentrated using a 10,000 molecular weight cutoff filter, then flash frozen in liquid nitrogen and stored at −80 °C.

Recombinant human IMPDH1(514) wildtype and S477D proteins were purified as described previously (Plana-Bonamaisó et al., 2020; Thomas et al., 2012). Briefly, BL21 (DE3) *E. coli* cells were transformed with pET15b constructs and expression was induced at room temperature (RT) for 12-14 h with 1 mM IPTG. Cell pellets were resuspended in binding buffer (50 mM Tris pH 8.0, 100 mM KCl, 30 mM imidazole, 1.5 M urea, 10 mM 2-mercaptoethanol) containing protease inhibitors (1 µg/mL leupeptin, 1 µg/mL pepstatin, 1 µg/mL antipain, 250 µM benzamidine, and 3 mM AEBSF). Lysates were sonicated then clarified at 17,000 x g for 30 min at 4 °C. His-tagged protein was captured on talon or nickel-nitriloacetic acid affinity resin. Protein was eluted with binding buffer containing 250 mM imidazole, the His tag was removed, and protein was dialyzed into activity buffer (100 mM KCl, 100 mM Tris-HCl pH 8.0, 1 mM DTT) with 20% glycerol for storage.

### IMPDH1 activity assays

Aliquots of IMPDH1 protein were diluted in activity buffer (20 mM HEPES, 100 mM KCl, 1 mM DTT, pH 7.0) to 1 µM and incubated with 1 mM ATP, 1 mM IMP, 1 mM MgCl_2_ and varying concentrations of GTP for 30 min at 20 °C in 96-well ultraviolet transparent plates (Corning, 3679). 100 µL reactions were initiated by addition of 300 µM NAD^+^. NADH production was measured by optical absorbance at 340 nm at 25 °C once per minute for 15 min with a Varioskan Lux microplate reader (Thermo Fisher). Absorbance was correlated with NADH concentration using a standard curve. All data points are an average of three measurements from the same protein preparation. Error bars represent standard deviation.

### Negative stain electron microscopy

Protein samples (1 µM; 2.5 µL) diluted in assembly buffer (20 mM HEPES, 100 mM KCl, 1 mM DTT, pH 7.0) were applied to glow-discharged continuous carbon EM grids (Protochips) and stained with 0.7% uranyl formate as described previously (Johnson and Kollman, 2020). Grids were imaged by transmission EM at 100 kV accelerating voltage on an FEI Morgagni microscope equipped with a Gatan Orius CCD. Micrographs were collected at a nominal 22,000X magnification with a 3.9 Å pixel size.

### Cryo-EM sample preparation and data collection

Protein samples (1-5 µM; 2.5 µL) diluted in assembly buffer (20 mM HEPES, 100 mM KCl, 1 mM DTT, pH 7.0) were applied to glow-discharged C-flat holey carbon EM grids (Protochips), blotted at 4 °C with 100% relative humidity, and plunge-frozen in liquid ethane using a Vitrobot vitrification system (Thermo Fisher) as described previously (Johnson and Kollman, 2020). High-throughput data collection was performed with Leginon (Suloway et al., 2009) or SerialEM (Mastronarde, 2005) software packages controlling a Thermo Fisher Glacios TEM operating at 200 kV equipped with a Gatan K2 Summit direct electron detector or a Thermo Fisher Titan Krios G3 TEM at 300 kV equipped with a Gatan image filter and K3 Summit direct electron detector.

### Cryo-EM image processing

Movies were collected in super-resolution mode, then aligned and corrected for full-frame motion and sample deformation using the patch motion correction algorithm in CryoSPARC Live, with 2x Fourier binning and dose compensation applied during motion correction (Punjani et al., 2017). Contrast transfer function (CTF) estimation, initial particle picking, and two-dimensional (2D) classification were performed in CryoSPARC Live. CryoSPARC v3 or v4 was used for all subsequent image processing. Each dataset was processed individually using a similar workflow.

After several rounds of 2D classification to select the best-resolved classes, particles were boxed and re-extracted to generate a quality particle stack for further processing. Either *ab initio* reconstruction was used to generate an initial volume from these particles or a map from a previous IMPDH filament structure lowpass filtered to 30 Å was used as a starting volume for refinement. The D4 point-group symmetry of IMPDH filaments means there are two points along the filament that can be used as symmetry origins: the center of an octamer or at the assembly interface between octamers. Octamer-centered reconstructions of filaments were done with homogeneous or non-uniform refinement with D4 symmetry, per-particle defocus refinement, and per-group CTF refinement with a previously solved octamer-centered filament map (either extended or compressed) as a starting volume (Punjani et al., 2020). Interface-centered reconstructions were performed the same, except with a previously solved interface-centered filament map as a starting volume. Image processing workflows are provided for each structure (**Supp. Figs. 4, 5, 7**). Directional FSCs were calculated using the remote 3DFSC processing server (3dfsc.salk.edu) (Tan et al., 2017).

### Model building and refinement

Existing structures were used as templates for model building. Octamer-centered compressed IMPDH1(595)-WT (PDB: 7RGD) was a template for the IMPDH1(595)-S477D model. Octamer-centered compressed IMPDH1(546)-WT (PDB: 7RGQ), interface-centered compressed IMPDH1(546)-WT (PDB: 7RGI), octamer-centered extended IMPDH1(546)-WT (PDB: 7RGM), and interface-centered extended IMPDH1(546)-WT (PDB: 7RGL) were used as templates for IMPDH1(546)-S477D models. Initial templates were placed into cryo-EM densities by rigid-body fitting in UCSF ChimeraX (Goddard et al., 2018), then repeated cycles of semi-automated and manual fitting with ISOLDE (Croll, 2018) and automated fitting with Phenix real-space refinement (Liebschner et al., 2019) were used to generate final models. Ligand fits and other minor adjustments were also made with Coot (Emsley et al., 2010). Data collection parameters and refinement statistics are summarized in **Supp. Table 2**. Figures were prepared with UCSF ChimeraX.

### Human cell culture and transfection

HeLa cells (human cervical adenocarcinoma) were acquired from ATCC (#CCL-2) and maintained in DMEM medium (Gibco #10569010) supplemented with 10% fetal bovine serum (Gibco #10437028) in a 37 °C incubator set to 5% atmospheric CO_2_. Cells were subcultured 2-3 times per week with subcultivation ratios between 1:4 and 1:8 to maintain 50-80% confluence.

For transfection, HeLa cells were seeded at a density of 4 x 10^4^ cells per well in 8-well chambered glass culture slides (Celltreat #229168) and incubated overnight. The next day, culture medium was replaced with 345 µL fresh DMEM per well and mammalian expression plasmids were transfected into cells using the TransIT-HeLaMONSTER Transfection Kit (Mirus Bio #MIR2904) according to the manufacturer’s protocol. Briefly, transfections were prepared for each well as follows: 328 ng plasmid was mixed with 34 µL of Opti-MEM I Reduced Serum Medium (Thermo Fisher #31985070) prior to the addition of 1 µL TransIT-HeLa Reagent, gentle mixing, and then addition of 0.67 µL MONSTER Reagent and further mixing. This transfection mixture was incubated for 20 minutes at RT prior to dropwise addition to the appropriate well. Cells were incubated for 48 h without media change, then treated (or not) with 1 mM ribavirin (Sigma-Aldrich #R9644) for 2 h to induce filament assembly prior to fixation and immunofluorescence staining.

### Indirect immunofluorescence

HeLa cells were fixed with 4% paraformaldehyde for 15 min at room temperature (RT), washed 3x with PBS, then permeabilized with 0.1% Triton X-100 for 5 min at RT. Cells were washed 3x with PBS between all further steps. Cells were incubated with primary antibodies diluted in PBS for 1 h at RT, then with secondary antibodies diluted in PBS for 1 h at RT in the dark, and finally with 5 µM DAPI (Abcam ab228549) in PBS for 20 min at RT in the dark. The slide was mounted with ProLong Glass Antifade Mountant (Thermo Fisher #P36984) on #1.5 thickness coverglass (VWR #48393-251) and allowed to cure for 24 h prior to imaging. Slides were imaged on a Leica DM5500B widefield fluorescence microscope equipped with a Leica HCX PL Fluotar 40X 0.75 numerical aperture objective, Lumencor SOLA Light Engine solid-state light source, Leica DAPI ET, GFP ET, and TXR ET filters, and Hamamatsu ORCA-Flash 4.0 LT sCMOS camera controlled by Leica Application Suite X software.

Primary antibodies used: rabbit polyclonal IgG anti-IMPDH2 (1:500, Proteintech #12948-1-AP), mouse monoclonal IgG1 anti-FLAG epitope tag (1:200, Proteintech #66008-3-Ig). Secondary antibodies used: Alexa Fluor 488-conjugated goat anti-rabbit IgG (1:400, Abcam #ab150077), Alexa Fluor 594-conjugated goat anti-mouse IgG (1:400, Abcam #ab150120).

## DATA AVAILABILITY

Coordinates for atomic models are deposited in the PDB with accession codes: compressed IMPDH1(595)-S477D free octamer (**8U7M**); compressed IMPDH1(546)-S477D filament, octamer-centered (**8U7Q**); compressed IMPDH1(546)-S477D filament, interface-centered (**8U7V**); extended IMPDH1(546)-S477D filament, octamer-centered (**8U8O**); extended IMPDH1(546)-S477D filament, interface-centered (**8U8Y**). Cryo-EM maps are deposited in the Electron Microscopy Data Bank (EMDB) with accession codes: compressed IMPDH1(595)-S477D free octamer (**EMD-41986**); compressed IMPDH1(546)-S477D filament, octamer-centered (**EMD-41989**); compressed IMPDH1(546)-S477D filament, interface-centered (**EMD-42012**); extended IMPDH1(546)-S477D filament, octamer-centered (**EMD-42026**); extended IMPDH1(546)-S477D filament, interface-centered (**EMD-42029**).

## ACKNOWLEDGMENTS

The authors would like to thank the Arnold and Mabel Beckman Cryo-EM Center at the University of Washington for access to electron microscopes. We also thank Alex Merz for the use of a biosafety cabinet and incubators for human cell culture; Suzanne Hoppins and Andrea Wills for fluorescence microscope use; Kelli Hvorecny, Eric Lynch, and Lauren Salay for fruitful discussion. This work was supported by a Helen Hay Whitney Foundation postdoctoral fellowship to S.J.C.; the US National Institutes of Health grant nos. T32GM008268 to A.G.O. and A.L.B., F31EY030732 to A.L.B., R01GM118396, R35GM149542, R21EY031546, and S10OD032290 to J.M.K.; the Spanish Ministerio de Ciencia e Innovación-FEDER-Fondo Social Europeo grant PID2019-109671GB-I00 to R.M.B.

---

**Supplementary Figure 1.**
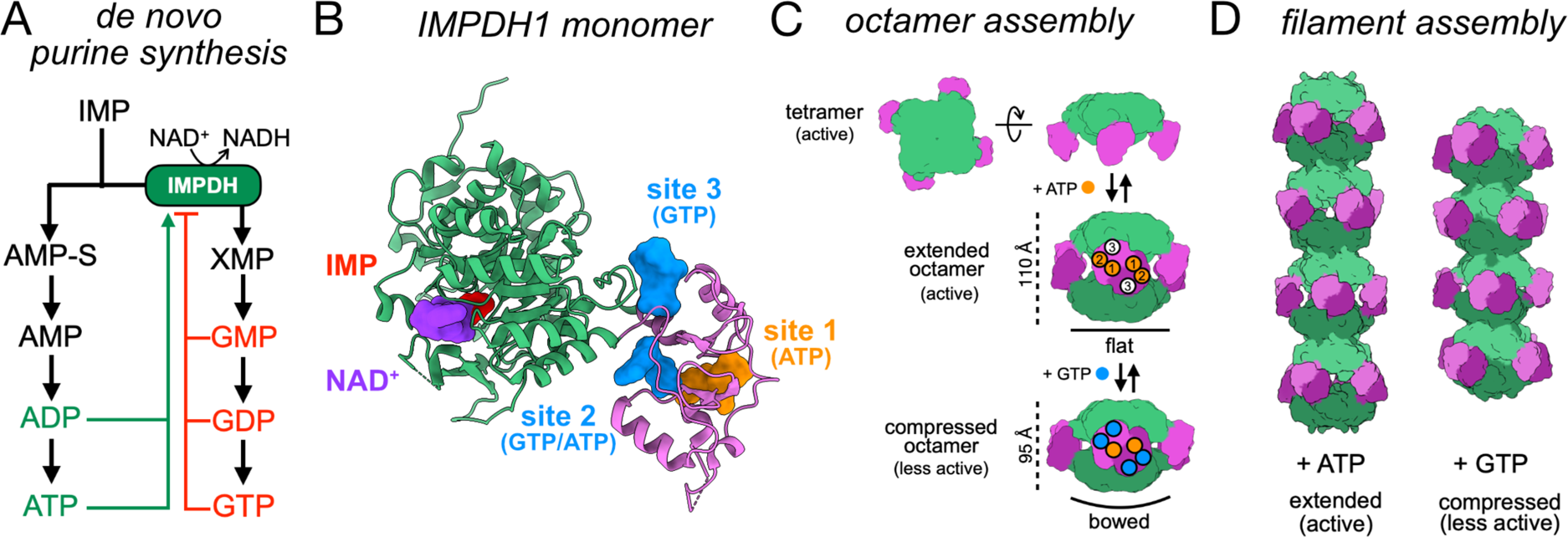
IMPDH function, structure, and regulation. **(A)** In the *de novo* purine synthesis pathway, IMP is converted to adenine or guanine nucleotides. IMPDH converts IMP to XMP in the rate-limiting step of GMP synthesis. Adenine and guanine nucleotides activate or inhibit IMPDH, respectively. **(B)** The IMPDH monomer (PDB: 7RGD) binds IMP (red) and NAD^+^ (purple) in the active site of the catalytic domain (green). ATP (orange) and GTP (blue) can bind in allosteric nucleotide binding sites in the Bateman domain (pink). **(C)** IMPDH is in equilibrium between tetramer and octamer in solution. Octamers form through interactions of the Bateman domains from opposing tetramers. The catalytic tetramer can adopt a flat or bowed conformation. Binding of nucleotides to the Bateman domain promotes either an extended, active or compressed, less active octamer conformation. **(D)** Octamers assemble end-on-end to form filaments that can accommodate both extended and compressed conformations.

**Supplementary Figure 2.**
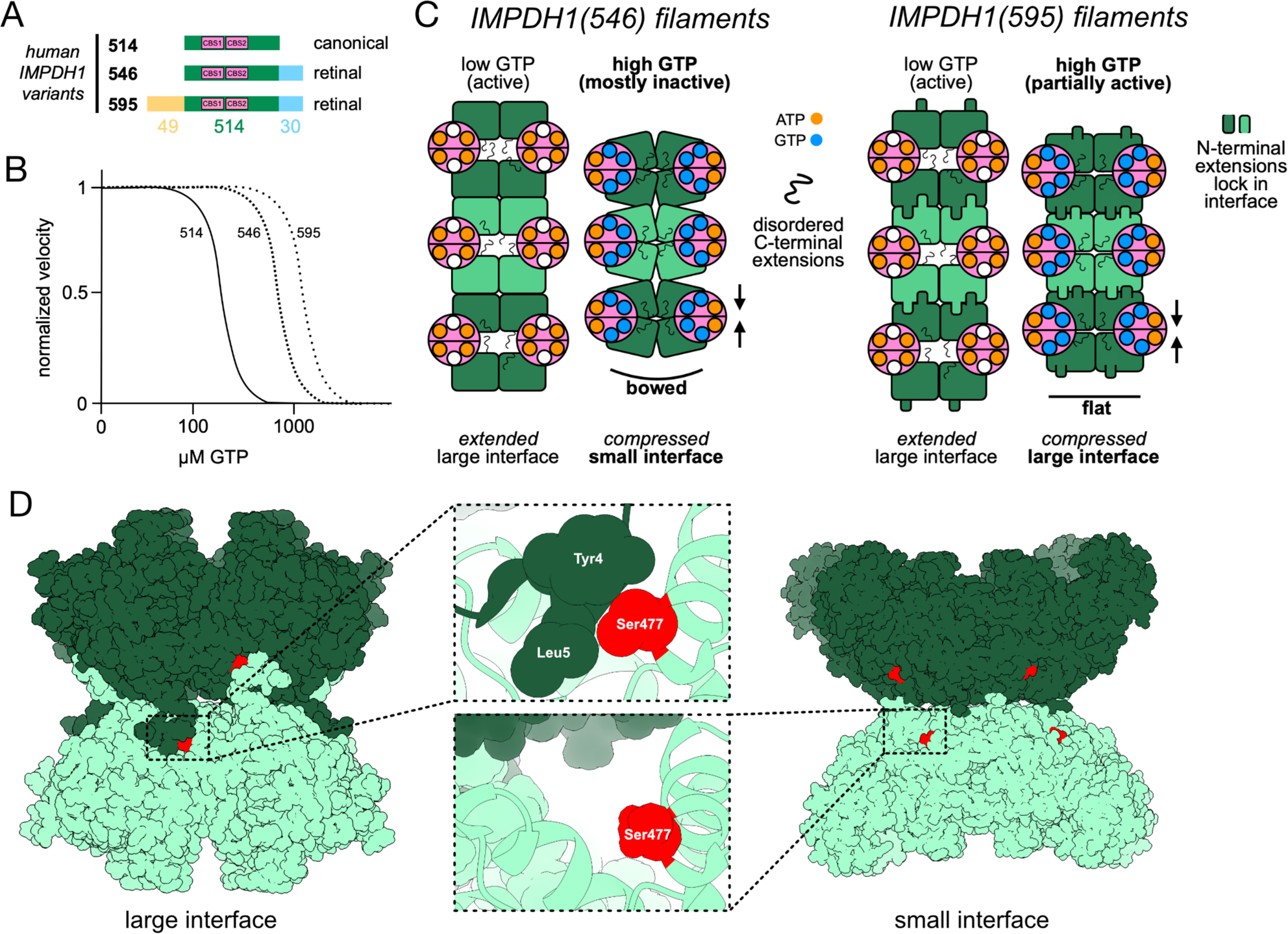
Human IMPDH1 retinal variants. **(A)** Schematic of human IMPDH1 variant structures. Pink: tandem CBS domains (Bateman domain). Blue: C-terminal extension. Gold: N-terminal extension. **(B)** Illustration of the relationship between IC_5_0 for GTP for the IMPDH1 variants. **(C)** Binding of ATP (orange) to IMPDH1(546) drives assembly of filaments containing extended octamers that are mostly active and connected with the large interface. GTP (blue) binding promotes a filament of less active, compressed octamers in a bowed conformation with the small interface. ATP also drives filament assembly of active, extended IMPDH1(595) octamers with the large interface. However, when GTP is present, the N-terminal extensions on the IMPDH1(595) filament maintain the flat conformation and large interface, resisting inhibition and remaining more active than IMPDH1(546). **(D)** Location of Ser477 (red) in both the large (PDB: 7RER) and small interfaces (PDB: 7RFE).

**Supplementary Figure 3.**
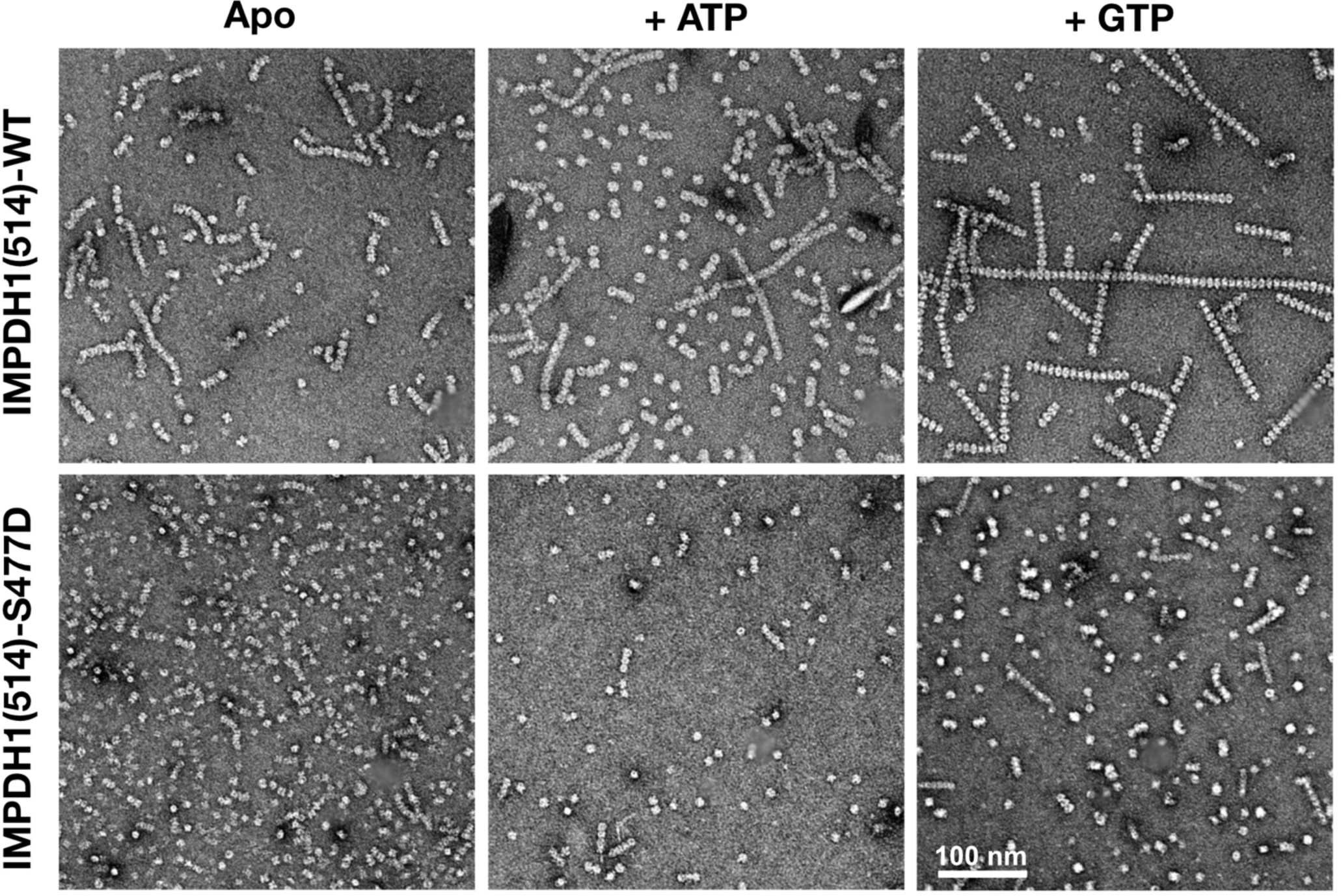
S477D partially disrupts IMPDH1(514) filament assembly. Negative stain EM of IMPDH1(514)-WT and IMPDH1(514)-S477D under Apo, + 1 mM ATP, or + 1 mM GTP conditions.

**Supplementary Figure 4.**
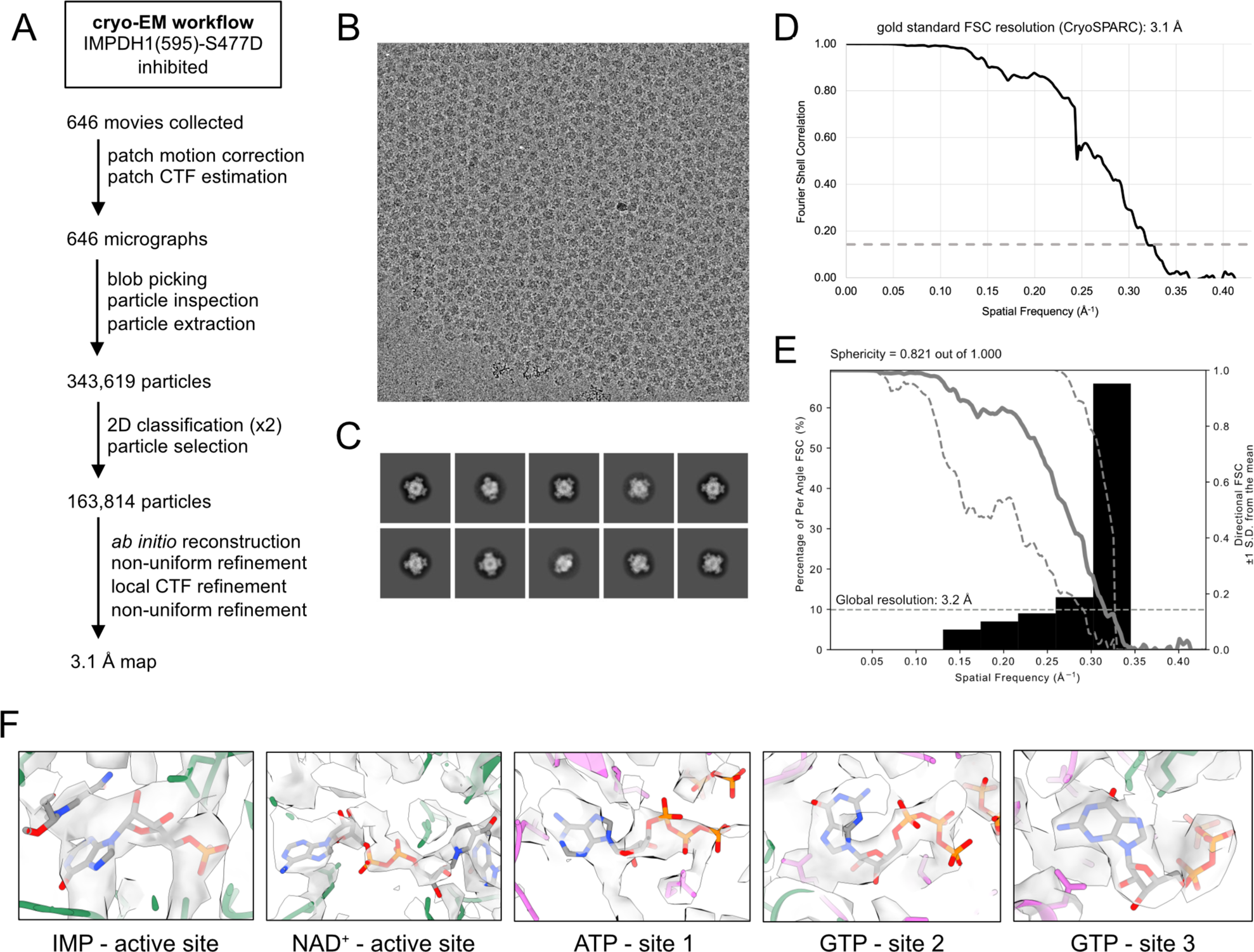
Cryo-EM workflow for the inhibited IMPDH1(595)-S477D structure. **(A)** Summarized data processing workflow. **(B)** Example micrograph. **(C)** Example 2D classes. **(D)** FSC curve from CryoSPARC. **(E)** Directional FSC plot depicting significant preferred orientation of the free octamer in ice. **(F)** Densities for bound ligands.

**Supplementary Figure 5.**
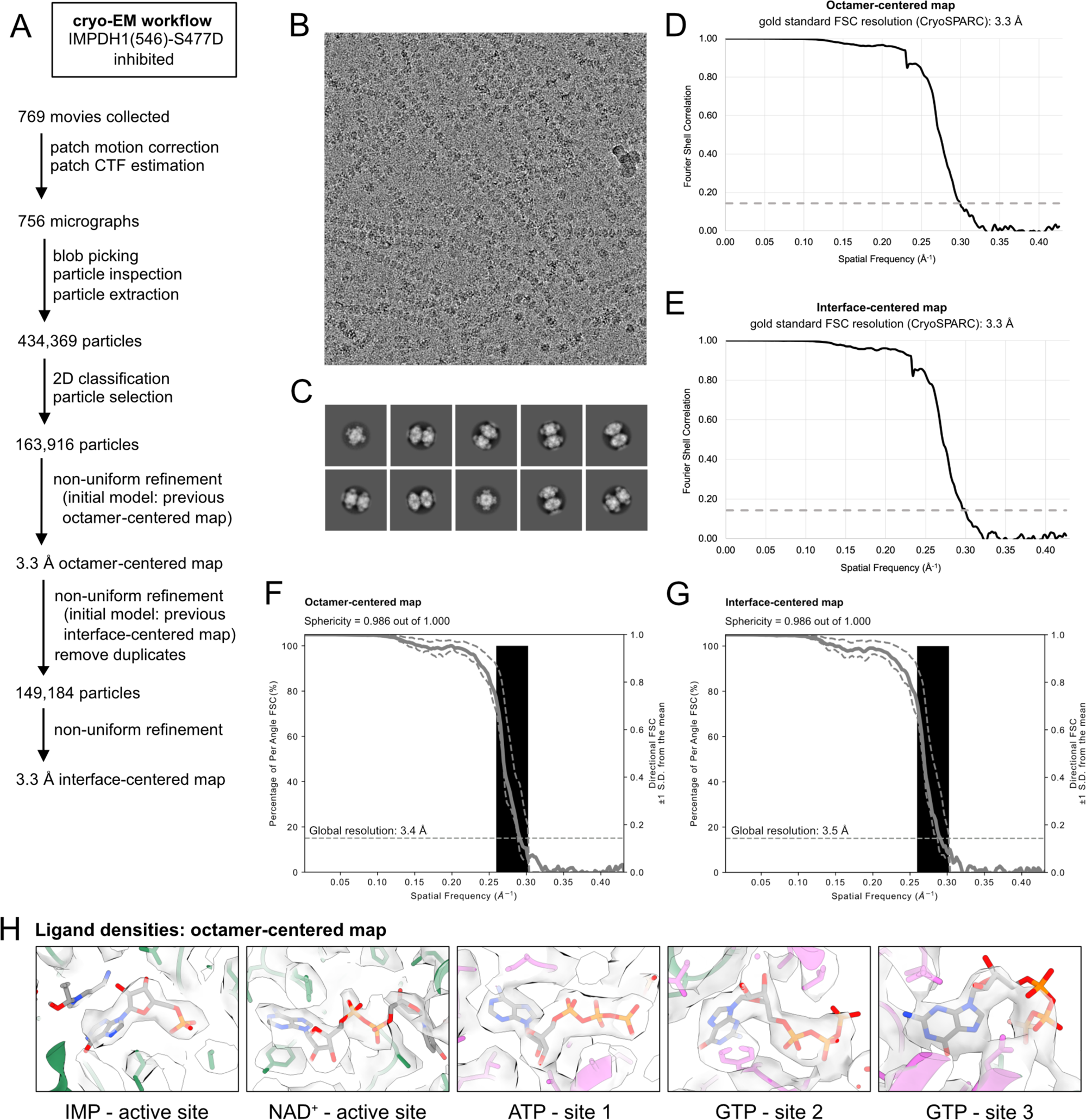
Cryo-EM workflow for the inhibited IMPDH1(546)-S477D structures. **(A)** Summarized data processing workflow. **(B)** Example micrograph. **(C)** Example 2D classes. **(D,E)** FSC curves from CryoSPARC for octamer-centered and interface-centered maps. **(F,G)** Directional FSC plots for octamer-centered and interface-centered maps. **(H)** Densities for bound ligands from the octamer-centered map.

**Supplementary Figure 6.**
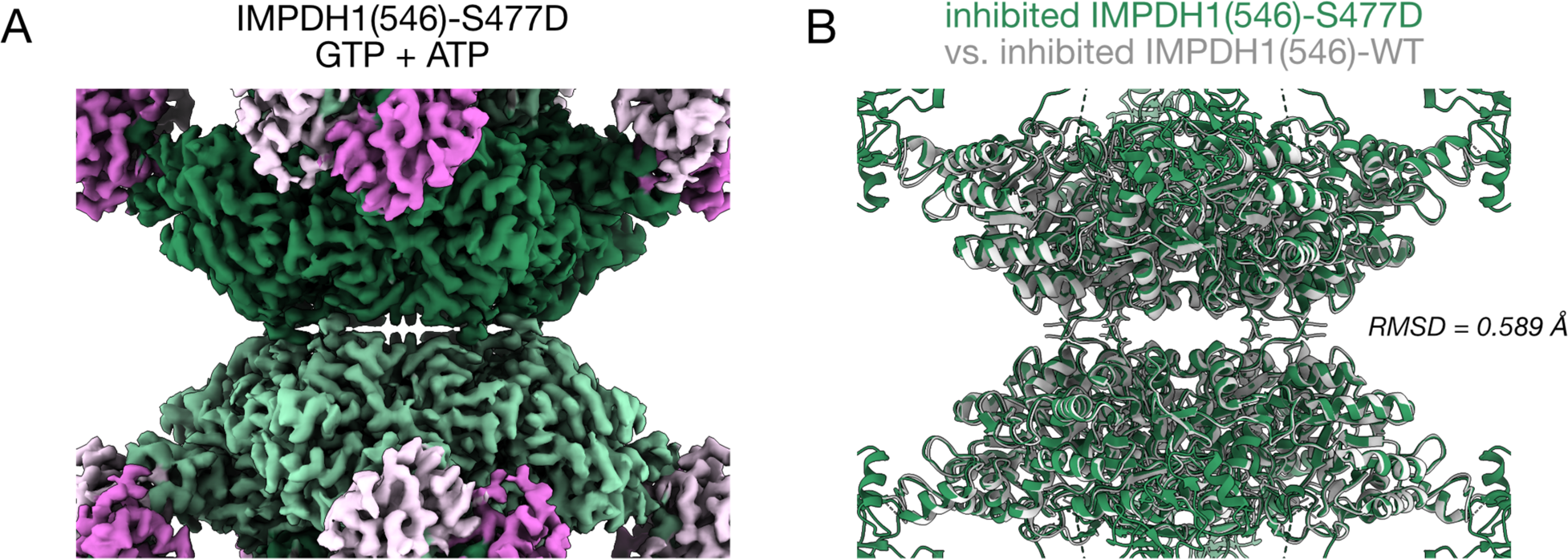
Interface-centered structure of inhibited IMPDH1(546)-S477D. **(A)** Cryo-EM map (3.3 Å) of the filament assembly interface between two compressed IMPDH1(546)-S477D octamers, in the presence of GTP, ATP, IMP, and NAD^+^. **(B)** Model of inhibited IMPDH1(546)-S477D (green) aligned to inhibited IMPDH1(546)-WT (grey). Aligned on chain A of both structures.

**Supplementary Figure 7.**
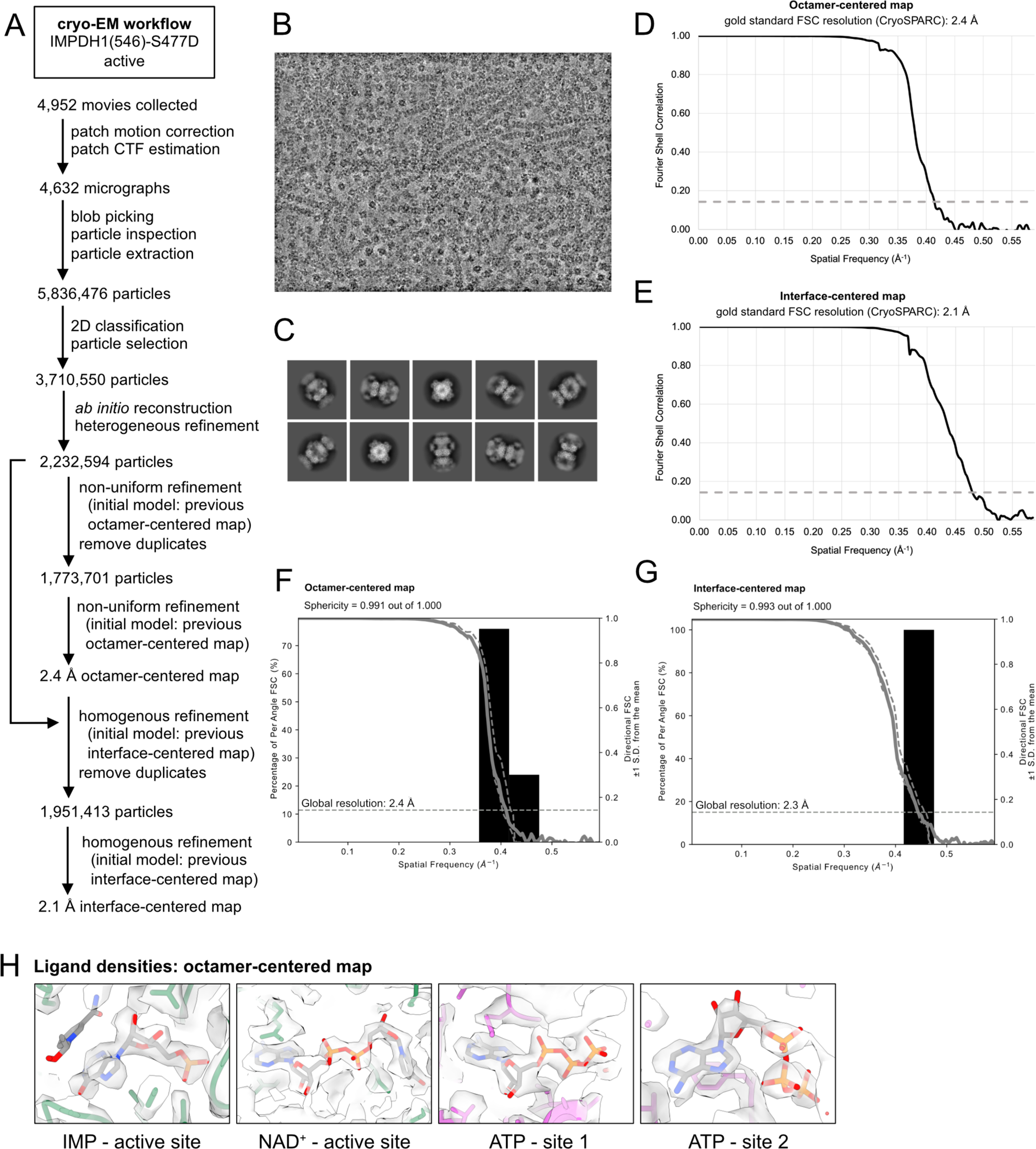
Cryo-EM workflow for the active IMPDH1(546)-S477D structures. **(A)** Summarized data processing workflow. **(B)** Example micrograph. **(C)** Example 2D classes. **(D,E)** FSC curves from CryoSPARC for octamer-centered and interface-centered maps. **(F,G)** Directional FSC plots for octamer-centered and interface-centered maps. **(H)** Densities for bound ligands from the octamer-centered map

**Supplementary Figure 8.**
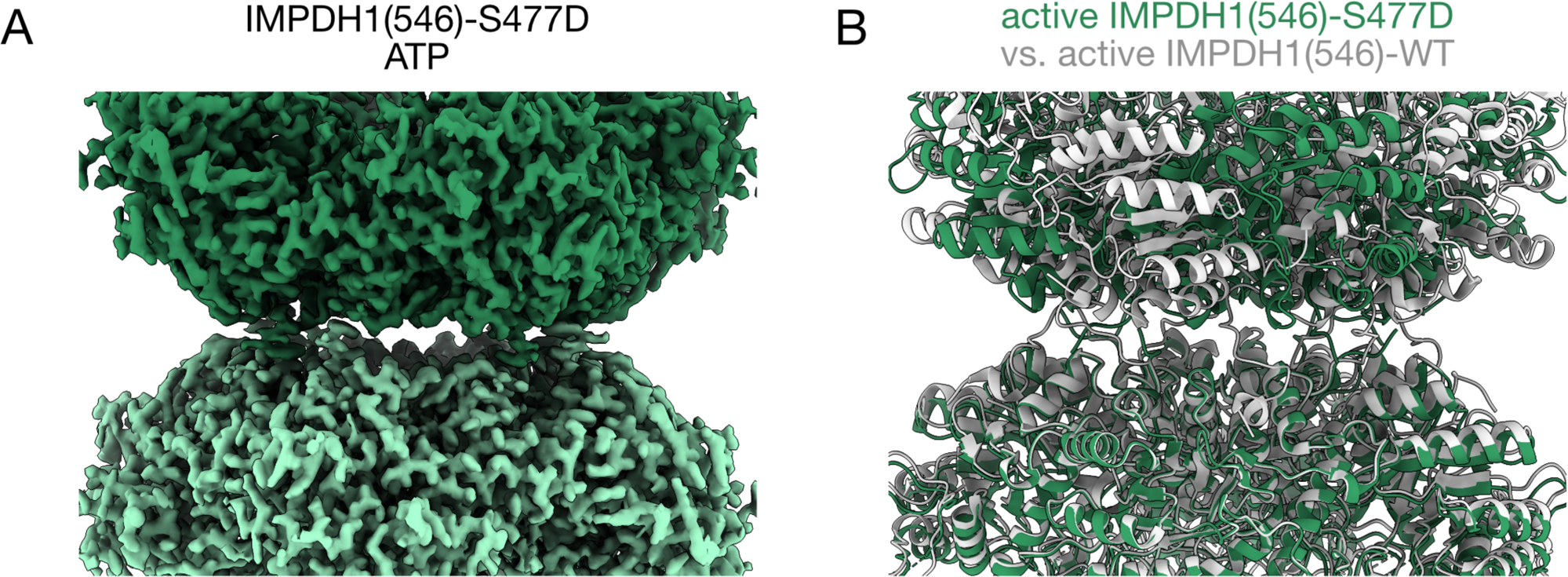
Interface-centered structure of active IMPDH1(546)-S477D. **(A)** Cryo-EM map (2.1 Å) of the filament assembly interface between two extended IMPDH1(546)-S477D octamers, in the presence of ATP, IMP, and NAD^+^. **(B)** Model of active IMPDH1(546)-S477D (green) aligned to active IMPDH1(546)-WT (grey). Aligned on chain A of both structures.

**Supplementary Figure 9.**
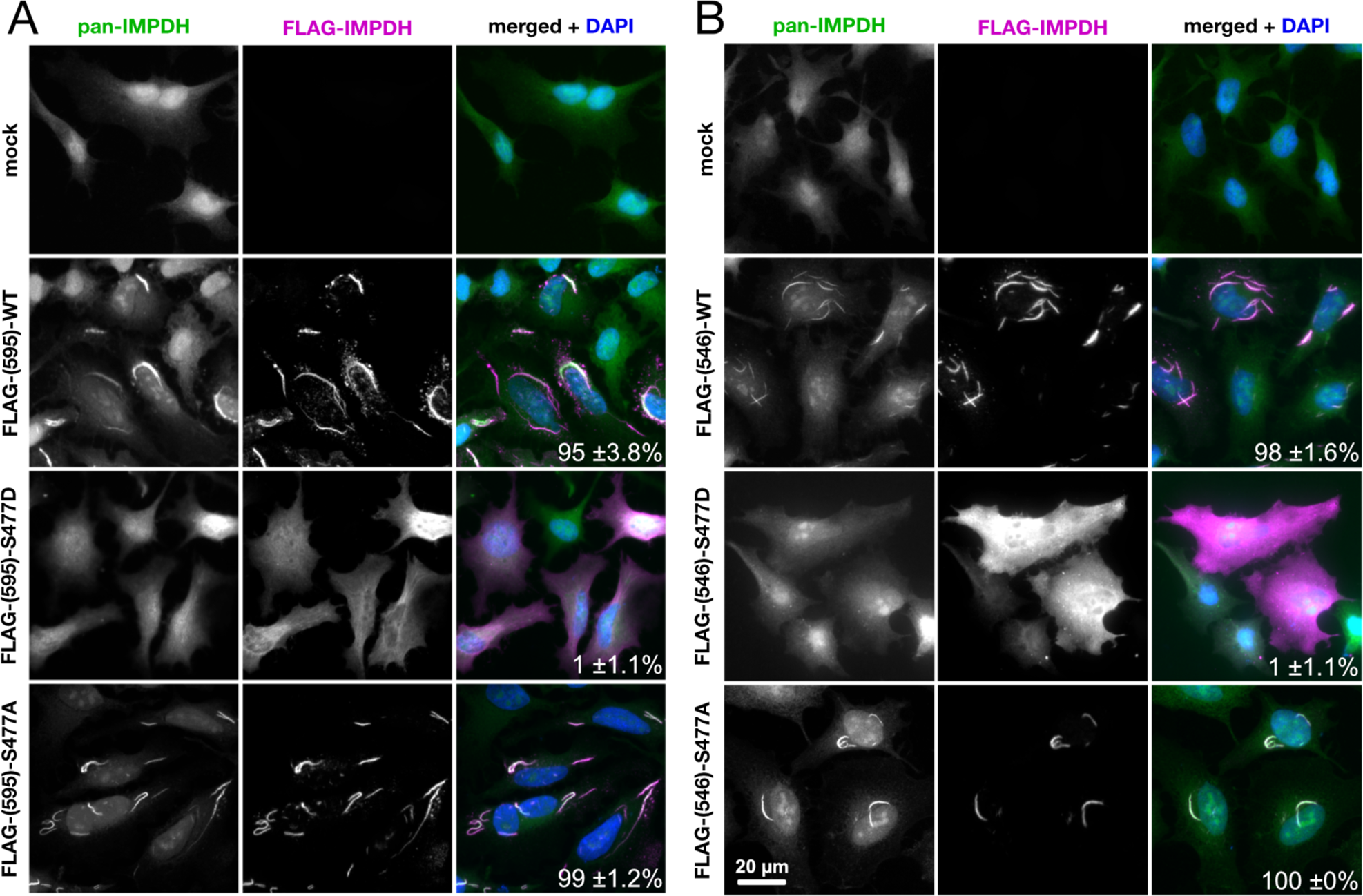
S477D disrupts filament assembly in cells without ribavirin treatment. **(A)** HeLa cells were transfected with FLAG-tagged IMPDH1(595)-WT, IMPDH1(595)-S477D, or IMPDH1(595)-S477A. Overexpression of the WT and S477A mutant induced filament assembly, but S477D did not. Mock transfection is shown as a control. Green: all IMPDH (endogenous + transfected). Magenta: FLAG-tagged fusion proteins. Blue: nuclei stained with DAPI. Percentage of transfected cells containing filaments is displayed in the bottom right corner of merged panels. **(B)** Same experiment as in panel A, but with FLAG-tagged IMPDH1(546)-WT, IMPDH1(546)-S477D, and IMPDH1(546)-S477A. Experiments were performed twice.

**Supplementary Table 1.**
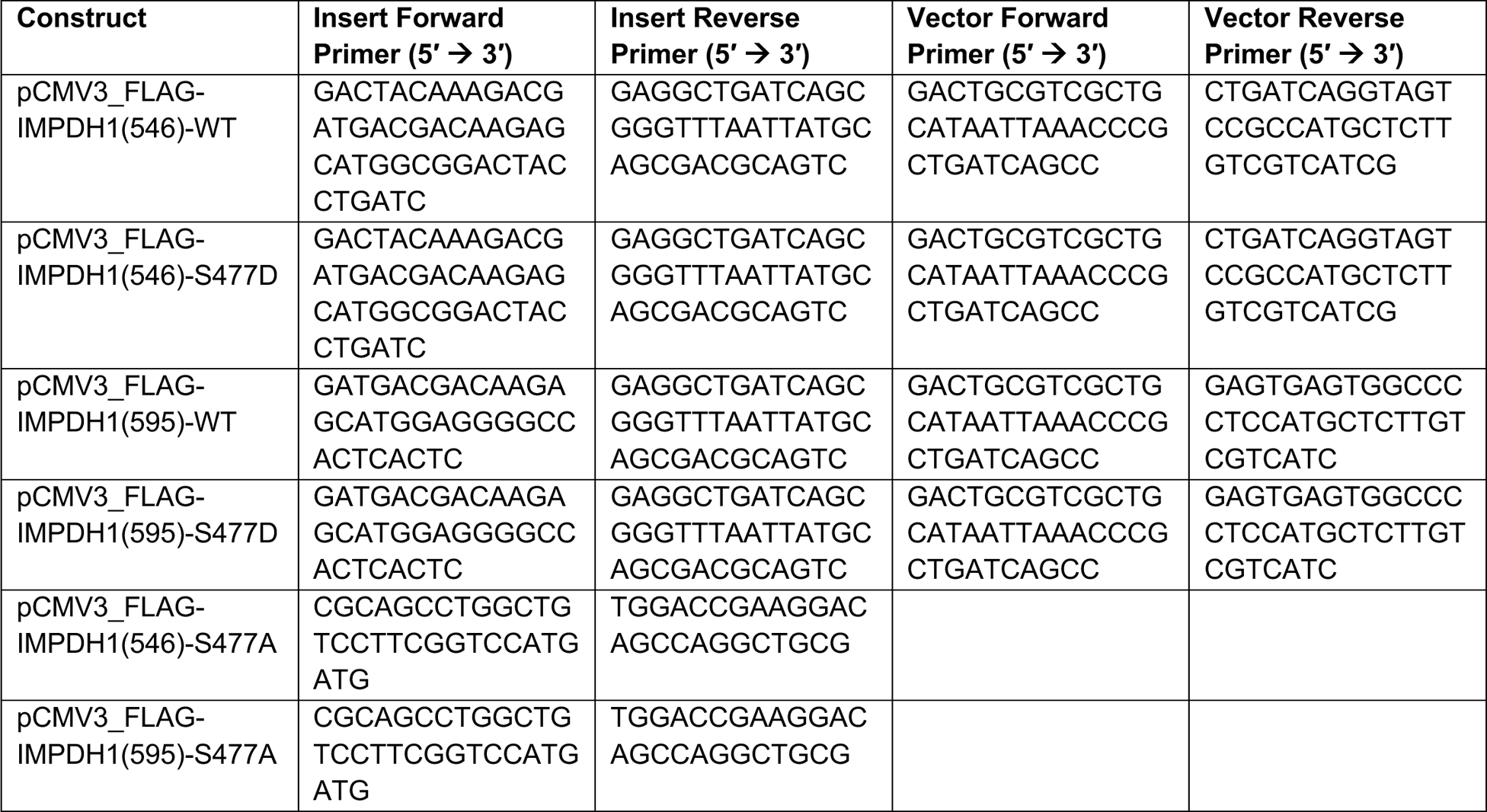
Primers used for cloning by Gibson Assembly.

**Supplementary Table 2.** Cryo-EM data collection, refinement, and validation statistics.

|  | IMPDH1(595)-<br>S477D<br>compressed<br>free<br>octamer | IMPDH1(546)-<br>S477D<br>compressed<br>filament<br>octamer | IMPDH1(546)-<br>S477D<br>compressed<br>filament<br>interface | IMPDH1(546)-<br>S477D<br>extended<br>filament<br>octamer | IMPDH1(546)-<br>S477D<br>extended<br>filament<br>interface |
| --- | --- | --- | --- | --- | --- |
| Ligands | GTP, ATP, IMP,<br>NAD<sup>+</sup> | GTP, ATP, IMP,<br>NAD<sup>+</sup> | GTP, ATP, IMP,<br>NAD<sup>+</sup> | ATP, IMP,<br>NAD<sup>+</sup> | ATP, IMP,<br>NAD<sup>+</sup> |
| PDB ID | 8U7M | 8U7Q | 8U7V | 8U8O | 8U8Y |
| EMDB ID | EMD-41986 | EMD-41989 | EMD-42012 | EMD-42026 | EMD-42029 |
| **Data collection and refinement** |  |  |  |  |  |
| Magnification | 36,000 | 36,000 | 36,000 | 105,000 | 105,000 |
| Voltage (kV) | 200 | 200 | 200 | 300 | 300 |
| Electron exposure (e<sup>-</sup>/Å<sup>2</sup>) | 65 | 50 | 50 | 50 | 50 |
| Defocus range (μm) | -1.8 to -1.2 | -1.8 to -1.2 | -1.8 to -1.2 | -1.8 to -1.2 | -1.8 to -1.2 |
| Pixel size (data collection) | 1.16 | 1.16 | 1.16 | 0.4215 | 0.4215 |
| Pixel size (reconstruction) | 1.16 | 1.16 | 1.16 | 0.843 | 0.843 |
| Micrographs (no.) | 646 | 756 | 756 | 4,632 | 4,632 |
| Initial particles (no.) | 343,619 | 434,369 | 434,369 | 5,836,476 | 5,836,476 |
| Final particles (no.) | 163,814 | 163,916 | 149,184 | 1,773,701 | 1,951,413 |
| Symmetry imposed | D4 | D4 | D4 | D4 | D4 |
| **Map resolution range (Å)** |  |  |  |  |  |
| Resolution (CryoSPARC) | 3.1 | 3.3 | 3.3 | 2.4 | 2.1 |
| Resolution (3DFSC) | 3.2 | 3.4 | 3.5 | 2.4 | 2.3 |
| **Model refinement and validation** |  |  |  |  |  |
| Initial model (PDB ID) | 7RGD | 7RGQ | 7RGI | 7RGM | 7RGL |
| **R.m.s. deviations** |  |  |  |  |  |
| Bond lengths (Å) | 0.017 | 0.017 | 0.017 | 0.014 | 0.011 |
| Bond angles (°) | 1.378 | 1.334 | 1.412 | 1.302 | 0.989 |
| MolProbity score | 1.79 | 1.60 | 1.66 | 1.44 | 1.30 |
| Clash score | 10.32 | 6.75 | 7.94 | 3.62 | 2.24 |
| C-beta deviations | 0% | 0% | 0% | 0% | 0% |
| Rotamer outliers (%) | 0.27% | 0.73% | 0.85% | 1.24% | 1.76% |
| **Ramachandran plot** |  |  |  |  |  |
| Favored (%) | 96.23% | 96.51% | 96.52% | 96.64% | 97.44% |
| Allowed (%) | 3.77% | 3.28% | 3.27% | 3.36% | 2.56% |
| Disallowed (%) | 0% | 0.21% | 0.22% | 0% | 0% |

